# The tilt board task as an internally valid practice-transfer paradigm for stabilometer balance assessments

**DOI:** 10.64898/2026.03.03.709278

**Authors:** Kimia Mahdaviani, Luc Tremblay, Alison Novak, Avril Mansfield

## Abstract

Practice–transfer paradigms are central to motor learning research, yet dynamic balance lacks standardized, internally valid practice–transfer task pairings. This study evaluated whether a mediolateral tilt board can serve as a valid transfer task for stabilometer-based balance assessment. Sixteen healthy young adults (20–35 years) completed a single session consisting of two 40-second trials on a mediolateral stabilometer and two 40-second trials on a mediolateral tilt board. Participants aimed to keep each platform horizontal during each trial. Performance outcomes were derived from platform deviation angle. Neuromuscular outcomes were derived from surface EMG of bilateral gluteus medius, vastus lateralis, and vastus medialis, including muscle synergy structure, bilateral co-activation index, RMS amplitude of muscle activation, and strategy ratios (hip-to-knee and asymmetry metrics). Between-task associations were assessed using Spearman correlations. Cross-task muscle synergy similarity was high (mean cosine similarity = 0.915 ± 0.044) and close to within-task trial-to-trial similarity, indicating preserved modular coordination across devices. Performance metrics were moderately to strongly correlated between tasks (RMS deviation angle: ρ = 0.621, p = 0.0089; time-in-balance: ρ = 0.668, p = 0.0036). EMG-derived strategy metrics also correlated significantly across tasks, including bilateral co-activation (ρ = 0.688, p = 0.0023), hip-to-knee ratio (ρ = 0.765, p = 0.0003), hip asymmetry ratio (ρ = 0.688, p=0.0023), and knee asymmetry ratio (ρ = 0.679, p = 0.0028). In contrast, EMG RMS amplitude did not correlate across tasks (ρ = −0.044, p = 0.873), suggesting task-specific gain of activation magnitude. Stabilometer and tilt board tasks shared a similar coordination structure and showed a high correlation in balance performance and neuromuscular strategy, supporting the tilt board as an internally valid transfer task for stabilometer-based dynamic balance paradigms. Similarity of tasks appears strongest at the level of modular control and strategy organization, with device-specific gain scaling of activation amplitude.

## INTRODUCTION

In motor learning research, transfer tasks are commonly used to determine whether the benefits of practice on one task generalize to other related tasks (Schmidt & Lee, 2011). Transfer is typically strong when the practiced and transfer tasks share key structural and informational features, a principle that can be referred to as the theory of identical elements (Aune et al., 2017). When movement goals, controlled variables, and sensory conditions align, performance improvements on a practiced task are more likely to carry over to an untrained one. Beyond mechanical similarity, theories of transfer-appropriate processing emphasize that overlap in perceptual and attentional demands also facilitates transfer (Lee, 1988; Roca et al., 2013). Together, these perspectives underscore the importance of carefully selecting transfer tasks that are distinct from the practiced task, yet internally aligned with it, i.e., sharing key controlled variables, task goals, and informational/sensory constraints (e.g., the relative reliance on proprioceptive/kinesthetic vs vestibular cues) that are hypothesized to support transfer (Bachman, 1961).

In many motor learning studies, the selection of practice and transfer tasks has become relatively standardized. For example, for gait training, where participants adapt to one split-belt walking condition and transfer is tested under a different treadmill walking context (e.g., altered belt-speed mapping, different walking direction, or other within-paradigm context changes; Hinkel-Lipsker & Hahn, 2017a, 2017b; Vasudevan et al., 2017). Similarly, in upper-limb motor control, practice often involves using a robotic arm under one force condition, with transfer evaluated by switching to a novel force condition (Dam et al., 2013; Krakauer et al., 1999, 2000; Shadmehr & Moussavi, 2000; Thoroughman & Shadmehr, 2000).

These designs are widely used because they provide standardized, controlled practice–transfer paradigms within established experimental families (e.g., locomotion on a treadmill under altered belt-speed mappings and upper-limb reaching with robotic force fields). However, there is a notable lack of standardized practice– transfer paradigms for dynamic balance tasks.

The primary objective of this study is to address this gap by exploring whether the stabilometer and tilt board balancing tasks can serve as a practice–transfer paradigm for dynamic balance control. To this end, we aim to determine whether performance on a mediolateral stabilometer correlates with performance on a mediolateral tilt board. While stabilometers and tilt boards are both used for assessing and training dynamic balance (Brouwer et al., 2019; Keller et al., 2023; Rizzato et al., 2025), the relationship between performance across them has not been systematically validated as a practice–transfer paradigm. This design aligns with motor learning theory by ensuring that both tasks share the same fundamental control structure while differing in mechanical configuration. If supported, this pairing could offer a within-laboratory internal validity standardized paradigm for investigating transfer in dynamic balance learning, similar to within-treadmill paradigms in locomotion and force-field adaptation paradigms in upper-limb control.

To achieve this, the present work will combine neuromuscular and behavioural outcomes to evaluate their correspondence across the two tasks. For neuromuscular outcomes, electromyography (EMG) features will be compared across tasks, as these features reflect stabilizing strategies and modular control patterns rather than a unitary sense of balance (Bachman, 1961; Torres-Oviedo & Ting, 2007). For behavioural outcomes, we will measure the correlation between stabilometer and tilt board balance performance using features derived from platform deviation angle that are widely used in balance-learning studies as indicators of balance performance (Giboin et al., 2015; McNevin et al., 2003; Shea & Wulf, 1999). By quantifying correlations and similarities across these domains, the study will provide a comprehensive validation of the stabilometer–tilt board pairing as a dynamic balance learning paradigm, bridging neuromuscular mechanisms with behavioural outcomes.

## MATERIALS AND METHODS

### Study design

This study employed a within-subjects design to investigate the relationship between performance on a mediolateral stabilometer (ST) and a mediolateral tilt board (TB). Each participant completed two trials on each device during a single experimental session.

### Participants

Sixteen healthy young adults (20–35 years) participated in this study. Participants were excluded if they had: difficulty understanding verbal or written English; or they reported any poor vision or poor hearing; any neurological or musculoskeletal condition; an injury that limits independent mobility; more than three months of training in dance, gymnastics, or other sporting or occupational activity with a large balance component; or participated in a moving platform study. The study was approved by the Research Ethics Board of the University Health Network (study ID: 21-6221), and participants provided written informed consent to participate. Participant age, sex, height, and mass were obtained.

### Apparatus

We used a custom made stabilometer, consisting of a swinging rectangular wooden platform (106 cm x 76 cm), which allows a maximum deviation of ± 30° degrees to either side of the horizontal plane of the platform. The tilt board was a rectangular platform (33 cm x 36 cm) with a cylindrical base, providing mediolateral instability. Both devices were placed on a non-slip surface. Figure 1 shows the stabilometer and tilt board used in this study along with their dynamics.

**Figure 1:**
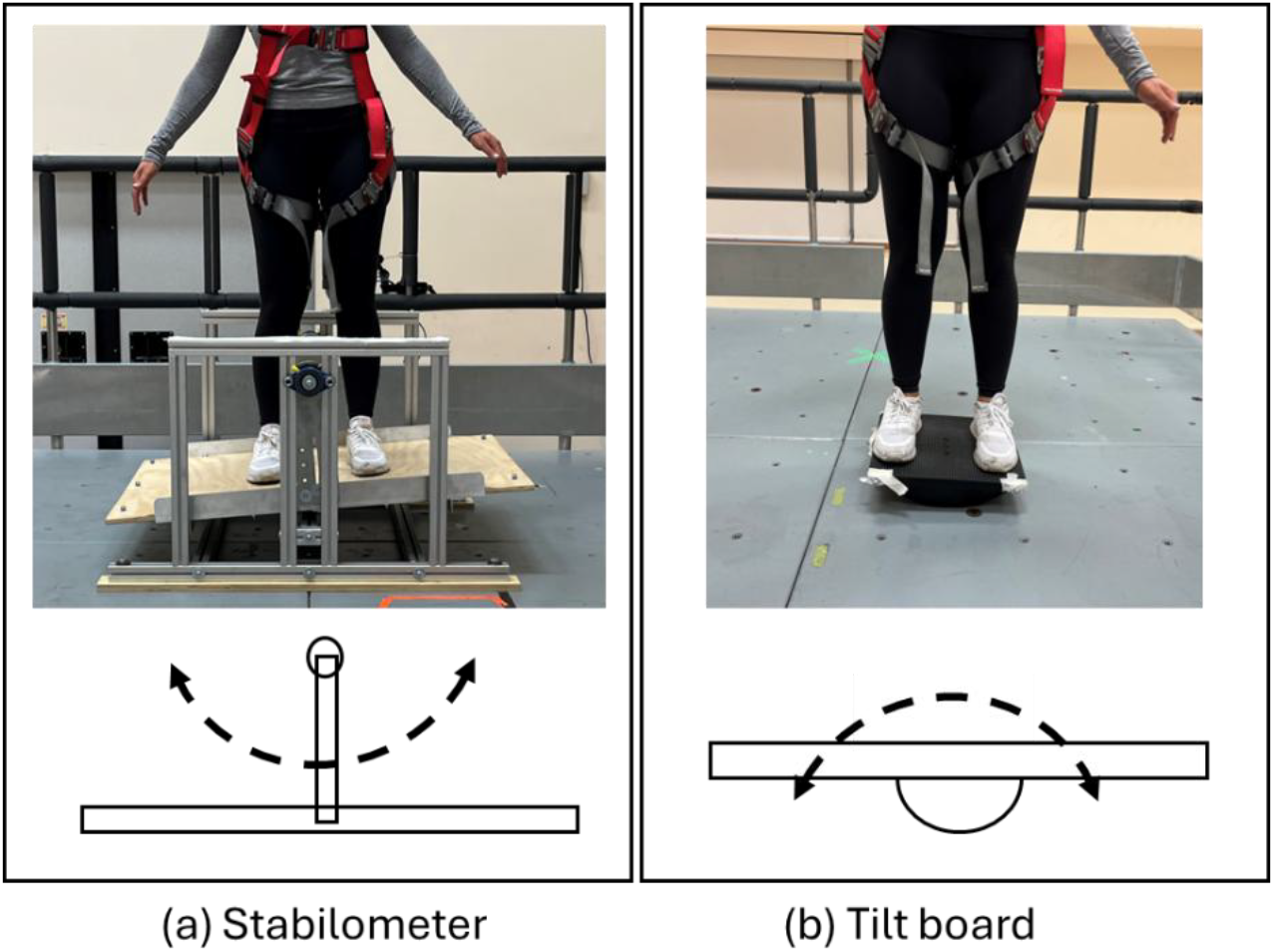
(a) Mediolateral Stabilometer, and (b) mediolateral tilt board, and their dynamic movement

Surface electromyography (EMG; Telemyo Direct Transmission System, Noraxon Inc., Scottsdale, Arizona, USA) was recorded from the Gluteus Medius, and Vastus Lateralis and Vastus Medialis of each leg during all trials with a sampling frequency of 1000 Hz. We used mildly abrasive exfoliating cream and alcohol wipes to clean patches of skin over the muscle bellies. Disposable silver/silver chloride electrodes were placed over the belly of each muscle. EMG data was then transmitted wirelessly from the connected sensors to the data collection computer via a receiver.

To track the movement of devices, we used motion capture system with nine motion capture cameras (Vicon Motion Systems Ltd, Oxford, UK) at 100 Hz and attached 5 reflective markers on the surface of each device. We extracted the trajectory of the attached markers and computed the tilt angles of both devices using trigonometry.

### Procedure

Experimental testing comprised two 45-second trials on each device where the first 5 seconds of each trial were removed from the measurements to account for stabilizing phase. Participants were instructed to keep the platform as close to horizontal as possible. Participants were asked to adopt the same stance for each task. Trials started when the device was in the horizontal position. The data collection session lasted approximately 1.5 hours, which allowed time for completing informed consent, instructing the participant, experimental trials, scheduled and requested rest breaks, setup and removal of equipment, and participant debriefing.

### Data processing

#### Neuromuscular outcomes

All data processing was conducted in MATLAB (R2023b, MathWorks, Natick, Massachusetts, USA). Preprocessing of EMG data followed a standardized pipeline. Raw signals were band-pass filtered (20–450 Hz, 4th-order Butterworth), detrended, full-wave rectified, and then low-pass filtered at 6 Hz to obtain the linear envelope (Fan et al., 2024; Konrad, 2005; McManus et al., 2020). Three primary EMG features were extracted; Muscle synergies were extracted using non-negative matrix factorization (NMF; Kubo et al., 2017; Lee & Seung, 1999; Ting & Macpherson, 2005). To this end, processed EMG signals of each trial were organized into a data matrix *X*, where each row corresponded to one muscle and each column represented a time sample. Muscle synergies were extracted using non-negative matrix factorization (NMF), which decomposes the EMG matrix into two non-negative matrices:

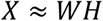

Here, *W* contains the synergy weight vectors, and *H* contains the corresponding temporal activation coefficients. Each column of *W* represents the relative contribution of each muscle to a given synergy, while each row of *H* describes the time-varying activation profile of that synergy. The number of synergies for each trial was defined as the smallest number of columns explaining at least 90% of total variance accounted for (VAF) and 80% of per-muscle VAF. To evaluate whether similar muscle coordination patterns were recruited across tasks, we quantified the correspondence between synergy weight vectors extracted independently from stabilometer balance task and tilt board balance task. Because the order of synergies generated by NMF is arbitrary, a matching procedure was implemented to establish direct correspondence between synergies derived from the two tasks. The synergy weight vectors from stabilometer balance task be denoted as *w*_*1,1*_, *w*_*1,2*_,*…,w*_*1,n*_, and those from tilt board balance task as *w*_*2,1*_, *w*_*2,2*_,*…, w*_*2,n*_. We first computed pairwise similarity values between all possible combinations of synergy vectors from the two tasks using a cosine similarity metric. This exhaustive comparison ensured that no assumptions were made regarding synergy order and allowed an unbiased assessment of cross-task similarity. For each synergy *w*_*1,i*_ from first task, we identified the synergy w_2,j_ from second task that produced the highest pairwise similarity value. These optimal matches were retained as the most representative correspondence between synergy structures across tasks. Once all synergies from first task were matched to their best counterpart in second task, an overall cross-task similarity index was computed by averaging the maximum similarity values across all *n* synergies:

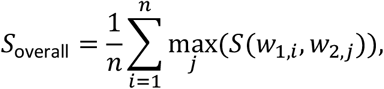

Where *S* is the cosine similarity between the two synergies. This aggregated similarity measure provided a single quantitative index describing the extent to which muscle coordination patterns were preserved across the two movement conditions. Higher overall similarity indicated greater reuse of common neuromuscular control strategies across tasks. As an upper-bound benchmark, we also computed the within-task (trial to trial) similarity for both stabilometer and tilt board.

Co-activation index (CAI) was calculated for bilateral muscle pairs as twice the minimum of paired activations divided by their sum, averaged across time and muscle pairs. RMS amplitude of EMG was computed from normalized envelopes and averaged across muscles. In addition to these classical features, strategy ratios were derived to quantify the relative contributions of hip vs. knee muscles and to detect left–right dominance in muscle recruitment. We averaged normalized envelopes of the vastus lateralis (VL), vastus medialis (VM), and gluteus medius (GM) bilaterally. A hip-to-knee ratio was computed as the summed activity of GM (bilateral) divided by the summed activity of VL and VM (bilateral), reflecting the relative use of hip versus knee musculature. We also quantified asymmetry as the normalized difference between left and right sides for the hip (GM) and knee (VL + VM) muscles, where positive values indicated greater left-side activity and negative values indicated greater right-side activity.

#### Performance outcomes

Two performance metrics were extracted from balance platform signals. Root-mean-squared (RMS) deviation angle and time-in-balance ratio. RMS deviation angle was computed as:

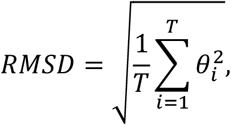

where *T* is the total trial time (in frames for a sampling rate of 100 frames per second) and *θ*_*i*_ is the deviation angle at frame *i*. time-in-balance ratio was defined as the cumulative duration that the platform remained within ±2.5° of horizontal, divided by the total duration of the trial.

### Data analysis

All analyses were conducted using SAS software (Version 9.4, SAS Institute, Cary, North Carolina, USA). For muscle synergy analyses, within-subject similarity values were calculated using cosine similarity. Associations between stabilometer and tilt board metrics were examined using Spearman correlation. For all scalar outcome measures (EMG-derived features and performance metrics), values were first computed separately for each trial and then averaged across the two trials within each task to obtain a single representative value per task per participant. For muscle synergy extraction, the two trials of each task were concatenated prior to non-negative matrix factorization. This procedure increases the robustness and stability of synergy estimation by providing a larger data sample for decomposition. To quantify within-task reliability, synergies were also extracted separately for each trial and compared using cosine similarity. To control the family-wise error rate across multiple related outcome measures, Holm–Bonferroni correction was applied within predefined families of tests (EMG-derived metrics and performance metrics). Statistical significance was set at p < 0.05.

## RESULTS

Participant characteristics of 16 participants are presented in Table 1.

**Table 1.** Participant characteristics.

|  | Mean or count | Standard deviation | Minimum | Maximum |
| --- | --- | --- | --- | --- |
| Age (years) | 24.5 | 3.66 | 20 | 32 |
| Sex (number) |  |  |  |  |
| Men | 8 |  |  |  |
| Women | 9 |  |  |  |
| Height (cm) | 170.1 | 9.8 | 155 | 183 |
| Body mass (kg) | 74.3 | 23.5 | 49.5 | 109.5 |

### Within-subject synergy similarity

The mean ± standard deviation of cross-task within-subject synergy similarity between the stabilometer and tilt-board tasks was 0.915 ± 0.044 with a median of 0.913 (cosine similarity range from 0 to 1). Within-task (trial to trial) similarity was 0.928 and 0.932 for stabilometer and tilt board tasks, respectively.

Figure 2 shows the histogram of within-subject muscle synergy similarity between the tasks, where for all participants this similarity is higher than 0.8.

**Figure 2:**
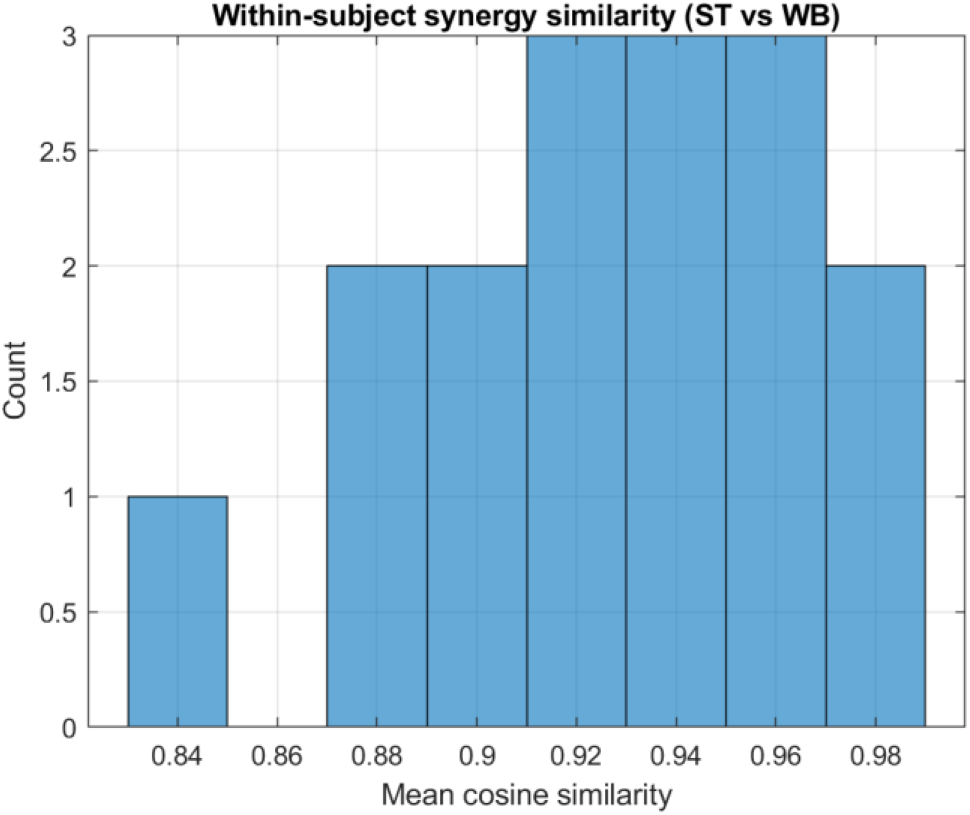
Histogram of within-subject muscle synergy similarity between Stabilometer (ST) and tilt board (TB) tasks. The height of each bin represents the number of participants with the synergy similarity in the bin range. Most subjects exhibited high similarity in synergy composition, with a mean cosine similarity of 0.915.

### Between-task correlations (EMG)

Between-task correlations for EMG-derived metrics are summarized in Table 2. The Co-activation Index (CAI) showed a strong and statistically significant positive correlation across tasks, with Spearman’s ρ = 0.688 (p = 0.0023), indicating consistent patterns of co-activation between tasks. The hip-to-knee ratio (ρ = 0.765, p = 0.0003), knee asymmetry (ρ = 0.679, p = 0.0028), and hip asymmetry (ρ = 0.688, p = 0.0023) all showed significant positive between-task correlations. In contrast, there was no correlation in RMS amplitude (Spearman’s ρ = −0.044, p = 0.873). All significant findings remained significant after correction.

**Table 2:** Between-task correlations (EMG). Values presented are Spearman correlation coefficients ρ with 95% confidence intervals in brackets, followed by the corresponding p-values. In addition, mean ± standard deviation values are provided for stabilometer and tilt board tasks.

| | Feature | Stabilometer | Tilt board | Spearman $\rho$ | p-value |
| --- | --- | --- | --- | --- | --- |
| EMG-derived metrics | Bilateral CAI | 0.680±0.068 | 0.681±0.102 | 0.688 [0.271, 0.878] | 0.0023 |
| | RMS amplitude | 0.126±0.063 | 0.130±0.065 | -0.044 [-0.527, 0.463] | 0.873 |
| | Hip/knee ratio | 0.137±0.088 | 0.180±0.149 | 0.765 [0.412, 0.910] | 0.0003 |
| | Hip asymmetry | -0.025±0.265 | -0.083±0.289 | 0.688 [0.271, 0.878] | 0.0023 |
| | Knee asymmetry | -0.048±0.159 | -0.071±0.154 | 0.679 [0.256, 0.874] | 0.0028 |
| Performance metrics | RMS dev angle (degree) | 7.6±2.5 | 9.0±3.9 | 0.621 [0.160, 0.848] | 0.0089 |
| | Time-In-Balance | 0.49±0.18 | 0.51±0.13 | 0.668 [0.236, 0.869] | 0.0036 |

### Between-task correlations (performance)

Between-task correlations of the performance on stabilometer and tilt board are given in Table 2. For RMS deviation angle of platform (ρ = 0.621, p = 0.0089) and time-in-balance ratio (ρ = 0.668, p = 0.0036), there were statistically significant positive correlations in outcomes between tasks. All significant findings remained significant after correction.

## DISCUSSION

The primary objective of this study was to validate the pairing of the mediolateral stabilometer (ST) and the mediolateral tilt board (TB) as a standardized practice-transfer paradigm for dynamic balance research. By employing a within-subjects design to assess both performance and neuromuscular control strategies, we aimed to determine the extent to which the central nervous system (CNS) generalizes motor solutions across mechanically distinct but functionally similar stability tasks. Performance, measured via RMS deviation angle and time-in-balance, demonstrated moderate-to-strong positive correlations, suggesting that skill in one task is predictive of, but not identical to, skill in the other. At the neuromuscular level, we observed a high similarity of muscle synergy structures and a strong correlation in co-activation strategies. However, the RMS amplitude of muscle activation did not correlate significantly between tasks. These findings suggest that while the CNS relies on the similar muscle synergies of motor modules and a consistent, individual-specific strategy for joint stiffness (co-activation) to do both tasks, the scaling of these outputs is highly sensitive to the specific mechanical impedance and sensory feedback loops of each device. This supports a hierarchical model of transfer, where high-level coordination policies are generalized, but low-level gain settings must be finely tuned to the specific physics of the environment (Diedrichsen & Kornysheva, 2015).

### Behavioural correspondence: the limits of transfer and task specificity

The moderate-to-strong behavioural correlations observed in this study sit at the intersection of two competing theories in motor learning: Thorndike’s classical Theory of Identical Elements (Woodworth & Thorndike, 1901) and the contemporary “specificity of balance” framework advocated by Giboin and colleagues (Giboin et al., 2015). Thorndike’s theory posits that transfer occurs to the extent that two tasks share common elements, whether they be physical properties or stimulus-response processing demands. The ST and TB tasks used in this study share significant structural elements: both require maintaining mediolateral equilibrium using similar lower body movements, and both engage vestibular and proprioceptive feedback loops to detect deviation from the horizontal. The high correlations found confirm that these shared elements facilitate a baseline level of positive transfer, validating the utility of performance on the stabilometer as a predictor for tilt board performance. However, the fact that performance correlations were not very high aligns with the “specificity of balance” hypothesis. This framework argues that balance is not a general ability but a set of highly specific skills optimized for the specific tasks (Giboin et al., 2018). Recent work by Rizzato et al. reinforced this by demonstrating that transfer of balance performance is highly dependent on the functional similarities between tasks, with improvements often failing to generalize even between mechanically similar unstable boards if the perturbation characteristic differs (Rizzato et al., 2025).

In our study, the lack of perfect performance correlation is likely the result of subtle but critical biomechanical discrepancies. The stabilometer operates on a fixed pivot (swinging dynamic), where the axis of rotation is static relative to the base. In contrast, the tilt board, with its cylindrical base, operates on a rolling pivot. This rolling mechanic introduces a translation of the centre of rotation and alters the lever arm of the ground reaction force vector relative to the ankle joint as the board tilts (Silva et al., 2018). As a result, the relationship between ankle movements and the vestibular signals is different for the two devices (Peterka, 2002). The moderate correlation likely indicates that participants used a generalized balance policy across tasks, but performance was modulated by differences in the underlying mechanical dynamics of each device.

### Neuromuscular modularity: conservation of muscle synergies

One of the most robust findings of this investigation is the high degree of similarity in muscle synergy structure between the ST and TB tasks. Importantly, cross-task similarity was only slightly lower than the within-task (trial-to-trial) similarity for each device, meaning that switching from the stabilometer to the tilt board altered synergy composition little beyond normal within-task variability. Because within-task similarity provides an empirical reliability ceiling for the synergy extraction procedure, this pattern suggests that the primary coordination modules were preserved across devices.

This finding provides strong empirical support for the theory of modular control, which posits that the CNS simplifies the control of redundant degrees of freedom by activating muscles in pre-programmed groups rather than individually (d’Avella et al., 2003). This concept, originally proposed by Bernstein (Bernstein, 1967) suggests that the CNS avoids the computational burden of controlling every muscle independently by using “synergies” as the fundamental building blocks of movement. The similarity of synergy structure across the ST and TB tasks suggests that the CNS relies on a shared set of motor primitives to generate corrective torque, despite differences in the mechanical behaviour of the two devices. This argument aligns with the work of Chvatal and Ting (Chvatal & Ting, 2013), who argued that the synergies used for reactive balance in standing are also recruited during walking. If synergies are robust enough to be shared across behaviours as distinct as standing and walking, it follows that they would be conserved across two standing balance tasks challenging the same anatomical plane. The neural economy of synergies implies that the skill of balancing on the stabilometer or tilt board does not require the creation of new muscle patterns. Instead, the CNS recruits existing mediolateral stability synergies, likely encoded in spinal or brainstem circuits (Bizzi & Cheung, 2013), and simply modulates their temporal recruitment. This supports the “modular control hypothesis” (Berger et al., 2013), which suggests that motor learning is achieved by fine-tuning how the CNS recombines a fixed set of muscle synergies, rather than by restructuring the synergies themselves. Recent computational modeling by Fukunishi et al. further suggests that the quality and stability of these modules are critical determinants of learning performance, reinforcing the idea that a robust synergy set is a prerequisite for effective transfer (Fukunishi et al., 2024).

### Co-activation: an individual trait and safety margin

The strong positive correlation in the Co-activation Index (CAI) across tasks indicates that the regulation of joint stiffness is a consistent, subject-specific strategy. Co-activation of left and right muscles increases the mechanical impedance of the joint, creating a safety margin that resists large deviations instantaneously, essentially stabilizing the limb before neural reflexes can intervene (Latash, 2018). Our results suggest that the level of co-activation is less a function of the specific device and more an intrinsic motor signature or trait of the individual. Some individuals adopt a high-impedance strategy to minimize risk in the face of uncertainty, while others rely on more efficient, phasic muscle bursts. The persistence of this trait across the ST and TB tasks reflects findings in stroke rehabilitation, where Banks et al. identified heterogeneity in co-contraction that was characteristic of the patient rather than the specific walking task (Banks et al., 2017). This has significant implications for the proposed practice-transfer paradigm. If a participant adopts a high-stiffness strategy on the stabilometer, they are highly likely to transfer this strategy to the tilt board. While this strategy is effective for preventing falls in early learning (cf. Bernstein’s freezing of degrees of freedom; Bernstein, 1967), persistent co-activation increases metabolic cost and joint loading. Therefore, the ST-TB paradigm may be particularly useful for identifying individuals with maladaptive high-stiffness strategies and training them to “free” their degrees of freedom in a controlled environment before transferring to more complex tasks. Furthermore, as shown by recent research on uncertainty in motor control, co-activation levels scale with environmental unexpected disturbances (Saliba et al., 2020); the consistency observed here suggests that participants perceived a similar level of threat or uncertainty from both devices, prompting a consistent impedance response (Burdet et al., 2001).

Importantly, Bernstein described skill acquisition as progressing through stages, with an initial phase characterized by ‘freezing’ degrees of freedom (reducing effective DOFs via stiffening/co-contraction) to stabilize performance, followed by later phases in which DOFs are gradually ‘released’ and coordination is reorganized as the skill becomes more efficient (Bernstein, 1967). In the present study, the relatively short practice duration may preferentially capture this early stabilization stage, which could help explain why participants relied on conservative coordination strategies that preserve synergy structure across tasks. Longer multi-day training may reveal later stages, release and reorganization, that could alter synergy recruitment and potentially the extent or form of transfer.

### Strategy ratios: proximal–distal control and lateralization

In addition to preserved synergy structure and co-activation, the strong cross-task correlation in the hip-to-knee ratio indicates that participants maintained a consistent proximal–distal control preference across devices. Classic postural-control models distinguish between ankle-dominant and hip-dominant strategies depending on perturbation and task demands (Horak & Nashner, 1986). Because mediolateral balance relies heavily on hip abductor (gluteus medius) contributions to pelvis and foot placement, a stable hip-to-knee weighting across tasks suggests that individuals exhibit a subject-specific effector distribution that remains consistent across mechanically distinct but functionally similar balance contexts (Roelker et al., 2019). The preserved hip and knee asymmetry further supports this interpretation, indicating a stable lateral recruitment bias across devices. Together, these findings indicate that transfer in this task pairing extends beyond performance metrics to include conserved patterns of effector use.

### Dissociation of amplitude and structure: the role of gain scaling

Despite the high correlations in synergy structure and co-activation strategies, there was no significant correlation in the RMS amplitude of muscle activation between the two tasks. This dissociation highlights the distinction between the qualitative organization of movement (coordination) and the quantitative scaling of movement (gain). The lack of RMS amplitude correlation is likely driven by the differing mechanical gains of the devices. Tilting on a stabilometer generates a restoring torque defined by the pendulum length and mass distribution, whereas the same tilt on a rolling tilt board generates torque defined by the radius of curvature and the shift in centre of pressure. To achieve the same outcome (horizontal stability), the CNS must scale the magnitude of the descending neural drive differently for each device. While practicing on the stabilometer can effectively train the coordination pattern required for the tilt board, it cannot perfectly train the force calibration. This suggests that transfer of learning in dynamic balance is hierarchical: coordination transfers readily, but gain scaling requires task-specific exposure to the error signals of the new environment.

### Limitations and future directions

The present study assessed cross-task transfer following short exposure within a single session. Motor learning literature indicates that coordination strategies evolve across stages of skill acquisition, transitioning from exploratory and co-contracted control toward more refined and efficient patterns with practice (Bernstein, 1967). It is therefore possible that the strategy ratios and synergy organization observed here reflect early-stage control characteristics rather than stable long-term motor solutions. For ecological validity, longitudinal training studies are needed to determine whether the strategies reported here remains stable, strengthens, or differentiates across devices with extended practice. Therefore, it remains to be seen if long-term training leads to a divergence in synergies as experts develop highly specialized solutions (Giboin et al., 2018).

This validation was conducted exclusively in young, healthy adults. While this controlled sample was appropriate for isolating task-related differences without confounding age- or pathology-related variability, it limits generalizability. Older adults and clinical populations often demonstrate altered co-activation, asymmetry, and proximal–distal balance strategies. Whether the preserved effector weighting and cross-task transfer observed here extend to populations with sensory, neuromuscular, or balance impairments remains unknown and warrants direct investigation.

## AUTHOR CONTRIBUTIONS

K.M., L.T., A.N., and A.M conceived and designed research; K.M. analyzed data; K.M. and A.M. performed experiments; K.M. interpreted results of experiments; K.M. prepared figures; K.M., L.T., A.N., and A.M. drafted manuscript; K.M., L.T., A.N., and A.M. edited and revised the manuscript; K.M., L.T., A.N., and A.M. approved final version of the manuscript;

## STATEMENTS AND DECLARATIONS

### ETHICAL CONSIDERATIONS

The study was conducted in accordance with ethical guidelines, received approval from Research Ethics Board of the University Health Network (study ID: 21-6221).

### CONSENT TO PARTICIPATE

All participants provided written informed consent before participation.

### CONSENT FOR PUBLICATION

This study does not include any identifiable personal data from individuals.

### DATA AVAILABILITY

Source data for this study are not publicly available due to privacy or ethical restrictions.

